# Genomic analysis of pathogenic isolates of *Vibrio cholerae* from eastern Democratic Republic of the Congo (2014-2017)

**DOI:** 10.1101/708735

**Authors:** Leonid M. Irenge, Jerôme Ambroise, Prudence N. Mitangala, Bertrand Bearzatto, Raphaël K.S. Kabangwa, Jean-François Durant, Jean-Luc Gala

## Abstract

**Background:** Over the past recent years, *Vibrio cholerae* has been associated with outbreaks in Sub Saharan Africa, notably in Democratic Republic of the Congo (DRC). This study aimed to determine the genetic relatedness of isolates responsible for cholera outbreaks in eastern DRC between 2014 and 2017, and their potential spread to bordering countries.

**Methods/Principal findings:** Phenotypic analysis and whole genome sequencing (WGS) were carried out on 78 clinical isolates of *V. cholerae* associated with cholera in eastern provinces of DRC between 2014 and 2017. SNP-based phylogenomic data show that most isolates (73/78) were *V. cholerae* O1 biotype El Tor with CTX-3 type prophage. They fell within the third transmission wave of the current seventh pandemic El Tor (7PET) lineage and were contained in the introduction event (T)10 in East Africa. These isolates clustered in two sub-clades corresponding to Multiple Locus Sequence Types (MLST) profiles ST69 and the newly assigned ST515, the latter displaying a higher genetic diversity. Both sub-clades showed a distinct geographic clustering, with ST69 isolates mostly restricted to Lake Tanganyika basin and phylogenetically related to *V. cholerae* isolates associated with cholera outbreaks in western Tanzania, whereas ST515 isolates were disseminated along the Albertine Rift and closely related to isolates in South Sudan, Uganda, Tanzania and Zambia. Other *V. cholerae* isolates (5/78) were non-O1/non-O139 without any CTX prophage and no phylogenetic relationship with already characterized non-O1/non-O139 isolates.

**Conclusions/Significance:** Current data confirm the association of both DRC O1 7PET (T)10 sub-clades ST69 and ST515 with recurrent outbreaks in eastern DRC and at regional level over the past 10 years. Interestingly, while ST69 is predominantly a locally endemic sequence type, ST515 became adaptable enough to expand across DRC neighboring countries.

**Author’s summary:** Cholera is a severe diarrheal disease caused by the Gram-negative bacterium *Vibrio cholerae.* After originating in Asia, the disease spread across sub-Saharan Africa, notably Democratic Republic of the Congo. The aim of our study was to assess the transmission pattern of *V. cholerae* strains prevailing in eastern DRC, and determine their genetic relatedness to strains from other African countries and other parts of the world. Between 2014 and 2017, we isolated *V. cholerae* from fecal samples of patients with acute diarrhea in eastern DRC, and subsequently examined the DNA of the bacteria. The results show that they all clustered in two genetic groups (ST69 and ST515) falling within the third transmission wave of the current seventh pandemic El Tor (7PET) lineage and T10 introduction event in East Africa. The genetic signature of ST515 may be involved in its adaptation to environmental conditions found in eastern DRC, and contribute to its extended geographic distribution. Indeed, unlike the locally endemic ST69, ST515 is spreading extensively through DRC cross-border countries such as South Sudan, Tanzania, Uganda and Zambia. This plainly justifies a regional strategy to strengthen the fight against cholera in eastern Africa.

## Introduction

Cholera is a life-threatening diarrheal disease caused by a Gram-negative comma-shaped bacterium called *V. cholerae* [1, 2]. Serogrouping based on the reactivity of antibodies with outer membrane lipopolysaccharide O-antigen has allowed defining more than 200 *V. cholerae*, among which only two (O1 and O139) are so far associated with epidemic or pandemic cholera [3]. Africa, a previously cholera-free continent [4], now bears the highest burden of the disease. Sub Saharan countries in particular have been the most affected and notably DRC, which now ranks in the world as one of countries most frequently affected by serious outbreaks [4–6]. Cholera has indeed become part of the DRC clinical landscape, with most cases reported in the eastern provinces along the Albertine Rift [7, 8]. The year 2017 has even experienced a dramatic expansion of the disease to new provinces in the center and west of the country, and of particular concern is the decreased susceptibility of *V. cholerae* to antimicrobial drugs in DRC [9].

During the current seventh cholera pandemic El Tor (7PET), at least three independent but temporally overlapping waves of global transmission have been identified by phylogenetic analyses in Africa [10–13], with at least 13 re-introduction events (T1-13) causing epidemics, each genetic lineage probably representing an independent introduction event to that location [4, 14]. Recent phylogenetic analysis of isolates associated with cholera outbreaks in DRC between 2006 and 2014 showed that all of them belonged to the 7PET, wave 3, T10 east African sub-lineage [4].

Understanding the dynamics of *V. cholerae* associated with recent cholera outbreaks in DRC is paramount in order to get insight into the mechanisms associated with the endemicity of the disease in the country, the epidemicity at local and regional level and the trace-back of infection sources. This study provides genomic information of *V. cholerae* isolates associated with cholera outbreaks, which occurred in eastern DRC between 2014 and 2017.

## Methods

### Ethical Considerations

Given the low level of literacy of the patients, rectal swabs were sampled with their oral informed consent. For children, the informed consent was obtained from their parent or guardian. This verbal consent was recorded, prior to sampling, by local first-line responders. Healthcare workers and physicians signed the following statement: “We have explained the study to the patient in the areas under investigation and are satisfied that he/she understands and consents to sampling”. Ethical approval to conduct the study was obtained from the Provincial Ministers of healthcare of North and South Kivu provinces (DRC192/CAB/MP-SASAFPP/NK/2018). The use of oral consent was approved by the Institutional Review Board of Université catholique de Louvain/ Saint-Luc academic Hospital.

### Study Design

Between 2014 and 2017, rectal swabs or stool were sampled from patients presenting with severe watery diarrhea and admitted in cholera treatment centers or other healthcare centers in North Kivu, South Kivu and Maniema provinces. Rectal swabs and stool samples were shipped in Carry Blair medium to the AMI-LABO laboratory in North-Kivu and the Centre de Diagnostic et de Recherche en Maladies in South-Kivu. Personal identifiers were removed so that analyses of stored isolates were not traceable to individual patients. Each sample was labeled using a code referring to the date and location of sample collection.

### Phenotype of *V. cholerae* clinical isolates

Samples were incubated in saline and alkaline peptone water broth during 6 hours and subsequently streaked onto thiosulfate-citrate-bile salts (TCBS) agar at 37°C for 16-24 hours. Large and flattened yellow colonies with opaque centers and translucent peripheries were sub-cultured on Luria-Bertani agar and subsequently characterized by phenotypic tests, i.e. microscopic examination, oxidase assay, and Kligler’s iron agar for fermentation of carbon hydrates. Isolates were further characterized by additional phenotype testing including Voges Proskauer assay (VP), hemolysis of sheep erythrocytes (HSE), chicken red cells agglutination (CCA) and susceptibility to polymyxin B (PB). *Enterobacter aerogenes* (ATCC13048) and *V. cholerae* O395 were used as positive and negative control for the VP assay respectively. Serotyping was carried out using the Polyvalent O1, Ogawa and Inaba antisera (Becton Dickinson, Erembodegem, Belgium) following the manufacturer’s recommendations.

### Antimicrobial Susceptibility Testing

The susceptibility to antimicrobial agents (i.e., ampicillin, doxycycline, erythromycin, nalidixic acid, chloramphenicol, ciprofloxacin, sulfamethoxazole-trimethoprim and tetracycline) was performed by the disk diffusion method. Susceptibility tests were interpreted using European Committee on Antimicrobial Susceptibility Testing (EUCAST) guidelines. *Escherichia coli* ATCC 35218 was used as a control for bacterial growth and susceptibility to antibiotic disks.

### Next-Generation Sequencing

Isolates were shipped to Belgium for whole genome sequencing and subsequent genomic analysis. *V. cholerae* isolates were cultured overnight in 10 ml Luria-Bertani broth. DNA was isolated using the phenol chloroform protocol [15]. DNA was quantified using the Nanodrop® and the Qubit® fluorometric quantitation (Thermo Fisher Scientific, Asse, Belgium) and normalized to 0.2 ng/µl. Genomic DNA was simultaneously fragmented and tagged with sequencing adapters in a single step using Nextera transposome (Nextera XT DNA Library Preparation Kit, Illumina, San Diego, CA, USA). Tagmented DNA was then amplified with a 12-cycle PCR, cleaned up with AMPure beads, and subsequently loaded on a MiSeq paired-end 2 x 150 (reagent kit V2 (300 cycles) or 2 x 300 bp (MiSeq reagent kit V3 (600 cycles) sequence run.

### Genomic analysis: genetic relatedness, toxin phage, drug resistance and virulence

Raw genomic data from each *V. cholerae* isolates were submitted to the European Nucleotide Archive (ENA, http://www.ebi.ac.uk/ena), and are available under accession number (ERP114722). In order to assess the genetic relatedness of DRC isolates with those from other African countries (e.g. Cameroon, Central African Republic, Kenya, Tanzania, Uganda, Zambia), Asia and South America, a large set of genomes, including the O1 El Tor N16961 and the pre-7^th^ pandemic O1 M66 isolates was downloaded from the European Nucleotide Archive (ENA), Genbank and Ensembl databases. Paired-end reads from each *V. cholera*e isolate were assembled *de novo* to construct a draft genome using the SPADES v.3.11.1 software [16]. The quality of *de novo* assemblies was assessed using the Quast software (version 4.5) [17]. Each draft genome was analyzed to identify the *V. cholerae* species-specific *ompW* [18], the O1 *rfb*V and O139 *wbf*Z serogroup-specific [19] as well as classical and El Tor biotype-specific (*ctxB*, *rstR* and *tcpA)* genes [20]. In addition, genomes were screened for the presence of the 7PET-specific gene VC2346 [21]. A SNP-based phylogenomic analysis was conducted using kSNP 3.0 for SNP identification and parsimony tree construction based on the core genome. A first tree included all DRC O1 7PET isolates and representative of 7PET isolates from all regions of the world. The Dendroscope v.3.5.9 was used to root the tree with the N16961 strain [22] as an outgroup. The next tree included non-O1/non-O139 isolates form DRC (n = 5) and from other countries (n = 11), as well as O1 representatives from 6th pandemic (n = 2), Gulf Coast (n = 4), pre-7th pandemic (n = 4) and 7PET isolates (n = 2), along with the outgroup *Vibrio metoecus* (isolate 07 2435) used to root the tree. The MLST analysis was performed on each isolate by using the MLST scheme developed by Octavia et *al* [23]. The nucleotide sequences of a new allele of the *metE* gene, and new allelic combinations creating a novel sequence type (ST) were sent to the MLST database curator for allele and ST assignment. The CTX prophage harbored by O1 DRC isolates was compared to representatives of known CTX prophages [24–25].

Raw data from each *V. cholerae* DRC isolate were aligned to the complete genome of the O1 El Tor reference N16961. Each file was screened for the presence of large deletions. The Freebayes v1.0.2 software [26] was used to call variants from the reference genome. The complete list of mutations was annotated using the SNPeff v.4.3 software [27] and only mutations with a high or moderate impact (i.e. frameshift deletion, non-sense point mutation, missense, and inframe deletion) were selected.

Each draft genome was then screened for the presence of virulence genes from the Virulence Factors Database (VFDB), selecting those which were experimentally tested, and for the presence of pathogenic islands (PAI) previously associated with various sub-lineages within the 7^th^ pandemic, namely virulence factors including Vibrio pathogenicity islands (VPI-1, VPI-2, VSP-1, VSP-2, a novel variant of VSP-2 (the VSP-2 WASA (West African-South America) and WASA-I, as well as other virulence genes [7, 28, 29]. A gene was deemed present if it matched the reference sequence, i.e. minimal identity match of 95% with a minimal coverage of 80% of the gene sequence, as previously described [30]. Each draft genome was also screened for the presence of antimicrobial resistance (AMR) genes. The complete list of screened genes was drawn up from the MEGARes database (https://megares.meglab.org). In order to selectively identify AMR genes acquired through horizontal gene transfer, the list based on MEGARes data was restricted to genes that were also found in the ResFinder database (https://cge.cbs.dtu.dk/services/ResFinder/), using BLASTn. In addition SNP-based AMR determinants were identified using ARIBA v.2.12.0 [31] with a home-made database including the *par*C, *gyr*A, *gyr*B, *par*E and *qnr* genes.

## Results

### Phenotypic results

Phenotypic features of *V. cholerae isolates* (n=78) are shown in Table 1. Most isolates (93.6%; 73/78) agglutinated with the *V. cholerae* Polyvalent O1 antiserum and were serotyped as El Tor biotype (47: VP+, CCA+, HSE+, and PB-resistant; 25: VP-, CCA+, HSE+, and PB-resistant; 2: VP+, CCA+, HSE+ and PB susceptible). Of these, 39 and 34 agglutinated with the Ogawa and Inaba antisera, respectively. Five isolates did not agglutinate with the Polyvalent O1 antiserum, being therefore typed as *V. cholerae* non-O1. Most positive VP reactions consisted in a light pink coloration, contrasting with the intense bright pink coloration which classically characterizes *V. cholerae* O1 El Tor isolates. Irrespective of their biotype, all *V. cholerae* isolates displayed resistance to co-trimoxazole and nalidixic acid, whilst retaining susceptibility to tetracycline and chloramphenicol. Nine *V. cholerae* O1 isolates displayed decreased susceptibility to ciprofloxacin, whereas 9 O1 and 2 non-O1 isolates were resistant to ampicillin (Table 1).

**Table 1.** Phenotypical features and antimicrobial susceptibility patterns of *V. cholerae* isolates from eastern DRC

|  | Isolate<br>identification | Isola-<br>tion<br>year | Original<br>specimen | Locatity | Province | Country | Serotype | Oxidase | VP | CCA | HSE | PXB | AMP | CHL | NA | CIP | SXT |
| --- | --- | --- | --- | --- | --- | --- | --- | --- | --- | --- | --- | --- | --- | --- | --- | --- | --- |
| 1 | CTMA-1402 | 2017 | Stools | Minova | South-Kivu | DR Congo | Inaba | Pos | Pos | Pos | Pos | R | R | S | R | S | R |
| 2 | CTMA-1403 | 2015 | Stools | Goma | North-Kivu | DR Congo | Inaba | Pos | Pos | Pos | Neg | R | R | S | R | S | R |
| 3 | CTMA-1404 | 2015 | Stools | Goma | North-Kivu | DR Congo | Inaba | Pos | Pos | Pos | Pos | R | R | S | R | S | R |
| 4 | CTMA-1405 | 2015 | Stools | Goma | North-Kivu | DR Congo | Inaba | Pos | Pos | Pos | Pos | R | S | S | R | R | R |
| 5 | CTMA-1406 | 2015 | Stools | Goma | North-Kivu | DR Congo | Inaba | Pos | Pos | Pos | Pos | R | S | S | R | S | R |
| 6 | CTMA-1407 | 2015 | Stools | Goma | North-Kivu | DR Congo | Inaba | Pos | Pos | Pos | Pos | R | S | S | R | R | R |
| 7 | CTMA-1408 | 2015 | Stools | Goma | North-Kivu | DR Congo | Inaba | Pos | Pos | Pos | Pos | R | S | S | R | S | R |
| 8 | CTMA-1409 | 2015 | Stools | Goma | North-Kivu | DR Congo | Inaba | Pos | Pos | Pos | Pos | R | S | ND | R | ND | R |
| 9 | CTMA-1410 | 2015 | Stools | Goma | North-Kivu | DR Congo | Inaba | Pos | Pos | Pos | Pos | R | R | S | R | S | R |
| 10 | CTMA-1411 | 2015 | Stools | Goma | North-Kivu | DR Congo | ND | Pos | Pos | Pos | Pos | R | S | S | R | S | R |
| 11 | CTMA-1412 | 2015 | Stools | Goma | North-Kivu | DR Congo | Inaba | Pos | Pos | Pos | Pos | R | R | S | R | S | R |
| 12 | CTMA-1413 | 2015 | Stools | Goma | North-Kivu | DR Congo | Inaba | Pos | Pos | Pos | Pos | R | S | ND | R | ND | R |
| 13 | CTMA-1414 | 2015 | Stools | Goma | North-Kivu | DR Congo | Inaba | Pos | Pos | Pos | Pos | R | S | 23 | R | S | R |
| 14 | CTMA-1415 | 2015 | Stools | Goma | North-Kivu | DR Congo | Inaba | Pos | Pos | Pos | Pos | R | S | S | R | S | R |
| 15 | CTMA-1416 | 2015 | Stools | Goma | North-Kivu | DR Congo | Inaba | Pos | Neg | Pos | Neg | R | R | S | R | S | R |
| 16 | CTMA-1417 | 2015 | Stools | Fizi-Baraka | South-Kivu | DR Congo | Ogawa | Pos | Pos | Pos | Neg | R | S | S | R | S | R |
| 17 | CTMA-1418 | 2015 | Stools | Fizi-Baraka | South-Kivu | DR Congo | Ogawa | Pos | Pos | Pos | Pos | R | ND | ND | R | ND | R |
| 18 | CTMA-1419 | 2015 | Stools | Fizi-Baraka | South-Kivu | DR Congo | Ogawa | Pos | Pos | Pos | Pos | R | S | S | R | S | R |
| 19 | CTMA-1420 | 2015 | Stools | Fizi-Baraka | South-Kivu | DR Congo | Ogawa | Pos | Neg | Pos | Pos | R | S | S | R | R | R |
| 20 | CTMA-1421 | 2015 | Stools | Fizi-Baraka | South-Kivu | DR Congo | Inaba | Pos | Neg | Pos | Pos | R | S | S | R | S | R |
| 21 | CTMA-1422 | 2015 | Stools | Fizi-Baraka | South-Kivu | DR Congo | Ogawa | Pos | Pos | Pos | Pos | R | S | S | R | S | R |
| 22 | CTMA-1423 | 2015 | Stools | Minova | South-Kivu | DR Congo | Inaba | Pos | Pos | Pos | Pos | R | S | S | R | S | R |
| 23 | CTMA-1424 | 2016 | Stools | Fizi-Baraka | South-Kivu | DR Congo | Inaba | Pos | Pos | Pos | Pos | R | ND | ND | R | ND | R |
| 24 | CTMA-1425 | 2016 | Stools | Fizi-Baraka | South-Kivu | DR Congo | Inaba | Pos | Pos | Pos | Pos | R | ND | ND | R | ND | R |
| 25 | CTMA-1426 | 2016 | Stools | Fizi-Baraka | South-Kivu | DR Congo | Ogawa | Pos | Pos | Pos | Pos | R | ND | ND | R | ND | R |
| 26 | CTMA-1427 | 2016 | Stools | Fizi-Baraka | South-Kivu | DR Congo | Ogawa | Pos | Pos | Pos | Pos | R | S | S | R | S | R |
| 27 | CTMA-1428 | 2016 | Stools | Fizi-Baraka | South-Kivu | DR Congo | Inaba | Pos | Pos | Pos | Pos | R | S | S | R | S | R |
| 28 | CTMA-1429 | 2016 | Stools | Fizi-Baraka | South-Kivu | DR Congo | Ogawa | Pos | Pos | Pos | Pos | R | S | S | R | S | R |
| 29 | CTMA-1430 | 2016 | Stools | Uvira | South-Kivu | DR Congo | Ogawa | Pos | Neg | Pos | Pos | R | ND | ND | R | ND | R |
| 30 | CTMA-1431 | 2016 | Stools | Uvira | South-Kivu | DR Congo | Ogawa | Pos | Neg | Pos | Pos | R | S | S | R | R | R |
| 31 | CTMA-1432 | 2016 | Stools | Uvira | South-Kivu | DR Congo | Ogawa | Pos | Neg | Pos | Pos | R | S | S | R | S | R |
| 32 | CTMA-1433 | 2016 | Stools | Uvira | South-Kivu | DR Congo | Ogawa | Pos | Neg | Pos | Pos | R | S | S | R | S | R |
| 33 | CTMA-1434 | 2016 | Stools | Uvira | South-Kivu | DR Congo | Ogawa | Pos | Pos | Pos | Pos | R | ND | ND | R | ND | R |
| 34 | CTMA-1435 | 2016 | Stools | Uvira | South-Kivu | DR Congo | Ogawa | Pos | Pos | Pos | Pos | R | S | S | R | S | R |
| 35 | CTMA-1436 | 2016 | Stools | Uvira | South-Kivu | DR Congo | Ogawa | Pos | Pos | Pos | Pos | R | S | S | R | R | R |
| 36 | CTMA-1437 | 2015 | Stools | Fizi-Baraka | South-Kivu | DR Congo | Ogawa | Pos | Pos | Pos | Pos | R | S | S | R | S | R |
| 37 | CTMA-1438 | 2015 | Stools | Fizi-Baraka | South-Kivu | DR Congo | Ogawa | Pos | Pos | Pos | Pos | R | ND | ND | R | ND | R |
| 38 | CTMA-1439 | 2015 | Stools | Fizi-Baraka | South-Kivu | DR Congo | Ogawa | Pos | Pos | Pos | Pos | R | S | S | R | S | R |
| 39 | CTMA-1440 | 2015 | Stools | Goma | North-Kivu | DR Congo |  | Pos | Pos | Pos | Pos | R | R | S | R | S | R |
| 40 | CTMA-1442 | 2016 | Stools | Alimbongo | North-Kivu | DR Congo | Inaba | Pos | Pos | Pos | Pos | R | S | S | R | S | R |
| 41 | CTMA-1443 | 2016 | Stools | Kabambare | Maniema | DR Congo | Ogawa | Pos | Pos | Pos | Pos | R | S | S | R | R | R |
| 42 | CTMA-1444 | 2016 | Stools | Kabambare | Maniema | DR Congo | Ogawa | Pos | Pos | Pos | Pos | R | ND | ND | R | ND | R |
| 43 | CTMA-1445 | 2016 | Stools | Fizi-Baraka | South-Kivu | DR Congo | Ogawa | Pos | Pos | Pos | Pos | R | ND | ND | R | ND | R |
| 44 | CTMA-1446 | 2016 | Stools | Fizi-Baraka | South-Kivu | DR Congo | Ogawa | Pos | Pos | Pos | Pos | R | ND | ND | R | ND | R |
| 45 | CTMA-1447 | 2016 | Stools | Fizi-Baraka | South-Kivu | DR Congo | Ogawa | Pos | Pos | Pos | Pos | R | S | S | R | R | R |
| 46 | CTMA-1448 | 2016 | Stools | Kimbilulenge | South-Kivu | DR Congo | Ogawa | Pos | Pos | Pos | Pos | R | S | S | R | S | R |
| 47 | CTMA-1449 | 2016 | Stools | Kimbilulenge | South-Kivu | DR Congo | Ogawa | Pos | Pos | Pos | Pos | R | ND | ND | R | ND | R |
| 48 | CTMA-1450 | 2014 | Stools | Fizi-Baraka | South-Kivu | DR Congo | Inaba | Pos | Pos | Pos | Pos | R | S | S | R | S | R |
| 49 | CTMA-1451 | 2014 | Stools | Fizi-Baraka | South-Kivu | DR Congo | Inaba | Pos | Pos | Pos | Pos | R | S | S | R | S | R |
| 50 | CTMA-1452 | 2014 | Stools | Fizi-Baraka | South-Kivu | DR Congo | Ogawa | Pos | Pos | Pos | Pos | R | S | S | R | R | R |
| 51 | CTMA-1453 | 2014 | Stools | Alimbongo | North-Kivu | DR Congo | Inaba | Pos | Pos | Pos | Pos | R | ND | ND | R | ND | ND |
| 52 | CTMA-1454 | 2015 | Stools | Masisi | North-Kivu | DR Congo | Inaba | Pos | Pos | Pos | Pos | R | ND | ND | R | ND | ND |
| 53 | CTMA-1455 | 2014 | Stools | Fizi-Baraka | South-Kivu | DR Congo | Ogawa | Pos | Pos | Pos | Pos | R | S | S | R | S | R |
| 54 | CTMA-1456 | 2014 | Stools | Fizi-Baraka | South-Kivu | DR Congo | Ogawa | Pos | Pos | Pos | Pos | R | S | S | R | S | R |
| 55 | CTMA-1457 | 2015 | Stools | Goma | North-Kivu | DR Congo | ND | Pos | ND | ND | ND | R | S | S | R | S | R |
| 56 | CTMA-1458 | 2015 | Stools | Goma | North-Kivu | DR Congo | ND | Pos | ND | ND | ND | R | R | S | R | S | R |
| 57 | CTMA-1459 | 2015 | Stools | Goma | North-Kivu | DR Congo | ND | Pos | ND | ND | ND | R | S | S | R | S | R |
| 58 | CTMA-1460 | 2015 | Stools | Goma | North-Kivu | DR Congo | Inaba | Pos | Pos | Pos | Pos | R | S | S | R | S | R |
| 59 | CTMA-1461 | 2015 | Stools | Goma | North-Kivu | DR Congo | Inaba | Pos | ND | ND | ND | R | S | S | R | S | R |
| 60 | CTMA-1462 | 2014 | Stools | Uvira | South-Kivu | DR Congo | Ogawa | Pos | Neg | Pos | Pos | R | S | S | R | S | R |
| 61 | CTMA-1463 | 2014 | Stools | Uvira | South-Kivu | DR Congo | Ogawa | Pos | ND | ND | ND | R | ND | ND | R | ND | R |
| 62 | CTMA-1464 | 2014 | Stools | Uvira | South-Kivu | DR Congo | Ogawa | Pos | Neg | Pos | Pos | R | S | S | R | S | R |
| 63 | CTMA-1465 | 2014 | Stools | Uvira | South-Kivu | DR Congo | Ogawa | Pos | Pos | Pos | Pos | R | S | S | R | S | R |
| 64 | CTMA-1466 | 2014 | Stools | Uvira | South-Kivu | DR Congo | Ogawa | Pos | Pos | Pos | Pos | R | S | S | R | S | R |
| 65 | CTMA-1467 | 2014 | Stools | Uvira | South-Kivu | DR Congo | Ogawa | Pos | Pos | Pos | Pos | R | R | S | R | S | R |
| 66 | CTMA-1468 | 2014 | Stools | Fizi-Baraka | South-Kivu | DR Congo | Ogawa | Pos | Pos | Pos | Pos | R | R | S | R | R | R |
| 67 | CTMA-1469 | 2014 | Stools | Fizi-Baraka | South-Kivu | DR Congo | Ogawa | Pos | Pos | Pos | Pos | R | S | S | R | S | R |
| 68 | CTMA-1470 | 2014 | Stools | Goma | North-Kivu | DR Congo | Inaba | Pos | Pos | Pos | Pos | R | ND | ND | R | ND | R |
| 69 | CTMA-1471 | 2014 | Stools | Fizi-Baraka | South-Kivu | DR Congo | Ogawa | Pos | Pos | Pos | Pos | R | ND | ND | R | ND | R |
| 70 | CTMA-1472 | 2014 | Stools | Fizi-Baraka | South-Kivu | DR Congo | Ogawa | Pos | Pos | Pos | Pos | R | R | S | R | S | R |
| 71 | CTMA-1473 | 2014 | Stools | Goma | North-Kivu | DR Congo | Inaba | Pos | Neg | Pos | Pos | R | S | S | R | S | R |
| 72 | CTMA-1474 | 2014 | Stools | Fizi-Baraka | South-Kivu | DR Congo | Ogawa | Pos | Neg | Pos | Pos | R | S | S | R | S | R |
| 73 | CTMA-1475 | 2014 | Stools | Fizi-Baraka | South-Kivu | DR Congo | Inaba | Pos | Pos | Pos | Pos | R | ND | ND | R | ND | R |
| 74 | CTMA-1476 | 2015 | Stools | Goma | North-Kivu | DR Congo | Inaba | Pos | Pos | Pos | Pos | R | S | S | R | S | R |
| 75 | CTMA-1477 | 2015 | Stools | Goma | North-Kivu | DR Congo | Inaba | Pos | Pos | Pos | Pos | R | S | S | R | S | R |
| 76 | CTMA-1478 | 2017 | Stools | Minova | South-Kivu | DR Congo | Inaba | Pos | Pos | Pos | Pos | R | S | S | R | S | R |
| 77 | CTMA-1479 | 2015 | Stools | Goma | North-Kivu | DR Congo | Inaba | Pos | Pos | Pos | Pos | R | S | S | R | S | R |
| 78 | CTMA-1480 | 2015 | Stools | Goma | North-Kivu | DR Congo | Inaba | Pos | Pos | Pos | Pos | R | S | S | R | S | R |

VP: Voges-Proskauer assay; CCA: chicken cell agglutination; HSE: hemolysis of sheep erythrocytes; PXB: Polymyxin B; AMP: Ampicillin; CHL: Chloramphenicol;
CIP: Ciprofloxacin; SXT: Sulfamethoxazole-Trimethoprim. Pos: Positive; Neg: Negative; ND: Not done

### Genomic results

The average size of draft genome assemblies was 4.06 and 3.83 for O1 and non-O1/non-O139 isolates, respectively, with all N_50_ values larger than 50.000 (see supplementary file). The average G+C content was determined to be 47.5 %. All DRC O1 isolates (n=73) were *ompW+, RfbV+*, *wbfZ-, tcpA*^*El Tor*^, *rstR*^*El Tor*^, *rtxC* and VC2346+, which characterize 7PET O1 *V. cholerae* [11]. Five isolates were *omp*W+, *Rfb*V*-, wbf*Z*-*, tcpA-, *rst*R-, *rstx*C-, corresponding to *V. cholerae* non-O1/non-O139. The SNP-based phylogeny unambiguously confirmed that the all current O1 isolates were associated with the sub-lineage T10 (7PET wave 3 clade) recovered in East Africa (Fig. 1) [25, 26]. This was further strengthened by the observation that they all carried the CTX-3 type of phage associated with this sub-lineage [26]. In line with these findings, *V. cholerae* O1 eastern DRC isolates clustered closely in 2 distinct sub-clades containing two MLST profiles, i.e. ST69 (39 isolates), and a newly assigned ST515 (34 isolates) (Fig. 1).

**Fig 1.**
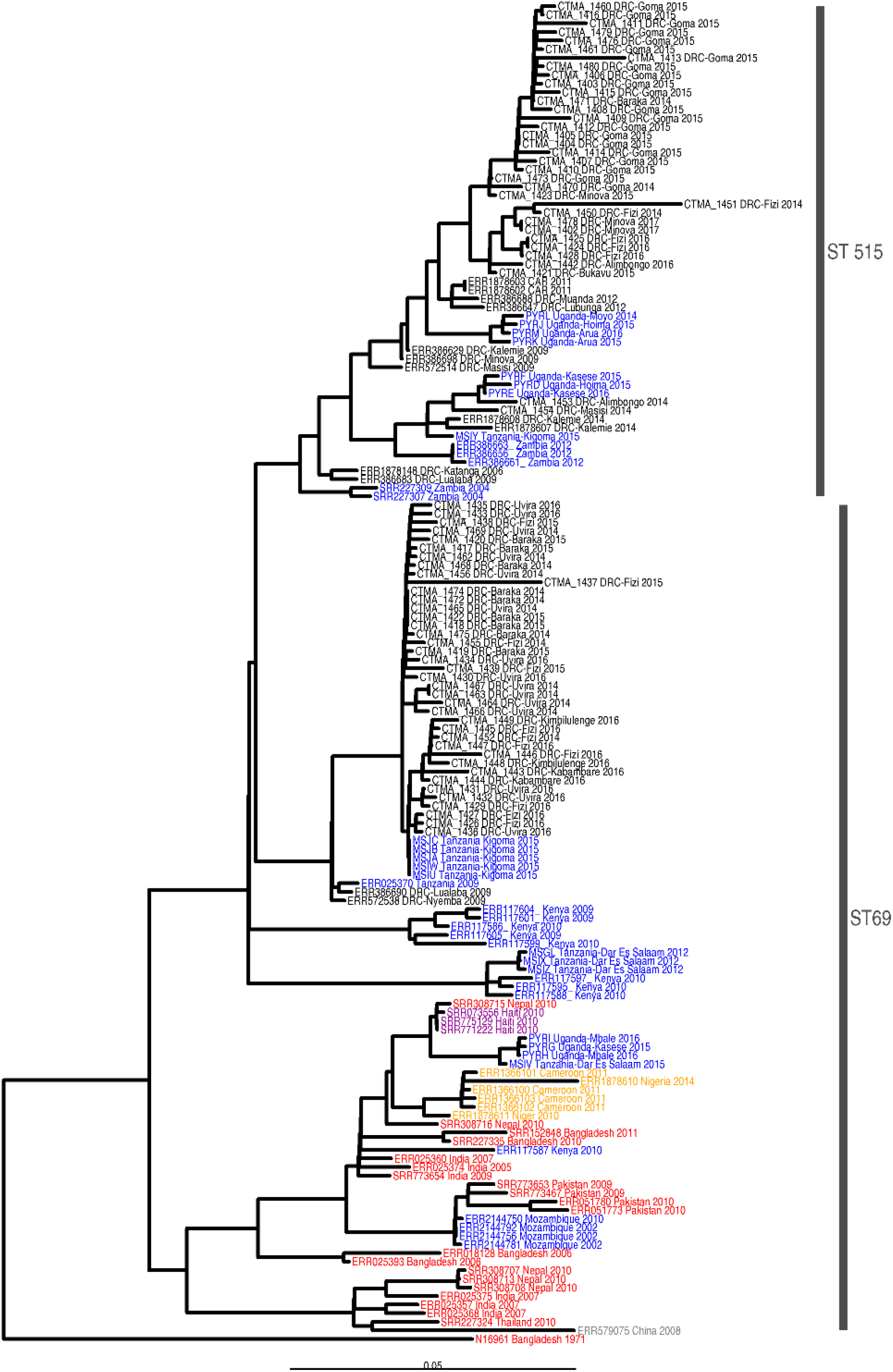
Phylogeny of seven pandemic (7PET) *V. cholerae* O1 isolates associated with cholera outbreaks in DR Congo between 2014 and 2017. The 7PET *V. cholerae* O1 biotype El Tor N19691 belonging to wave 1 was used as outgroup. The scale bar represents substitutions per variable site in the core genome. Green, blue, yellow, purple and red isolates represent 7PET wave 3 clades from Central Africa, East Africa, West Africa, Haiti and Asia regions.

Both sub-clades showed a distinct geographic pattern with ST69 sub-clade being found in the Lake Tanganyika basin (South-Kivu) and in Maniema provinces, and clustering together with 7PET *V. cholerae* isolates collected in Western Tanzania in 2015. While ST69 and ST515 were both identified in the Tanganyika basin, ST515 was the only sub-clade found in the Lakes Kivu and Edward basins and expanding northward (Fig. 2), hence covering a large area including three lake basins (Tanganyika, Kivu and Edward) and five bordering countries (DRC, Central African Republic, South-Sudan, Tanzania, Uganda and Zambia).

**Fig 2.**
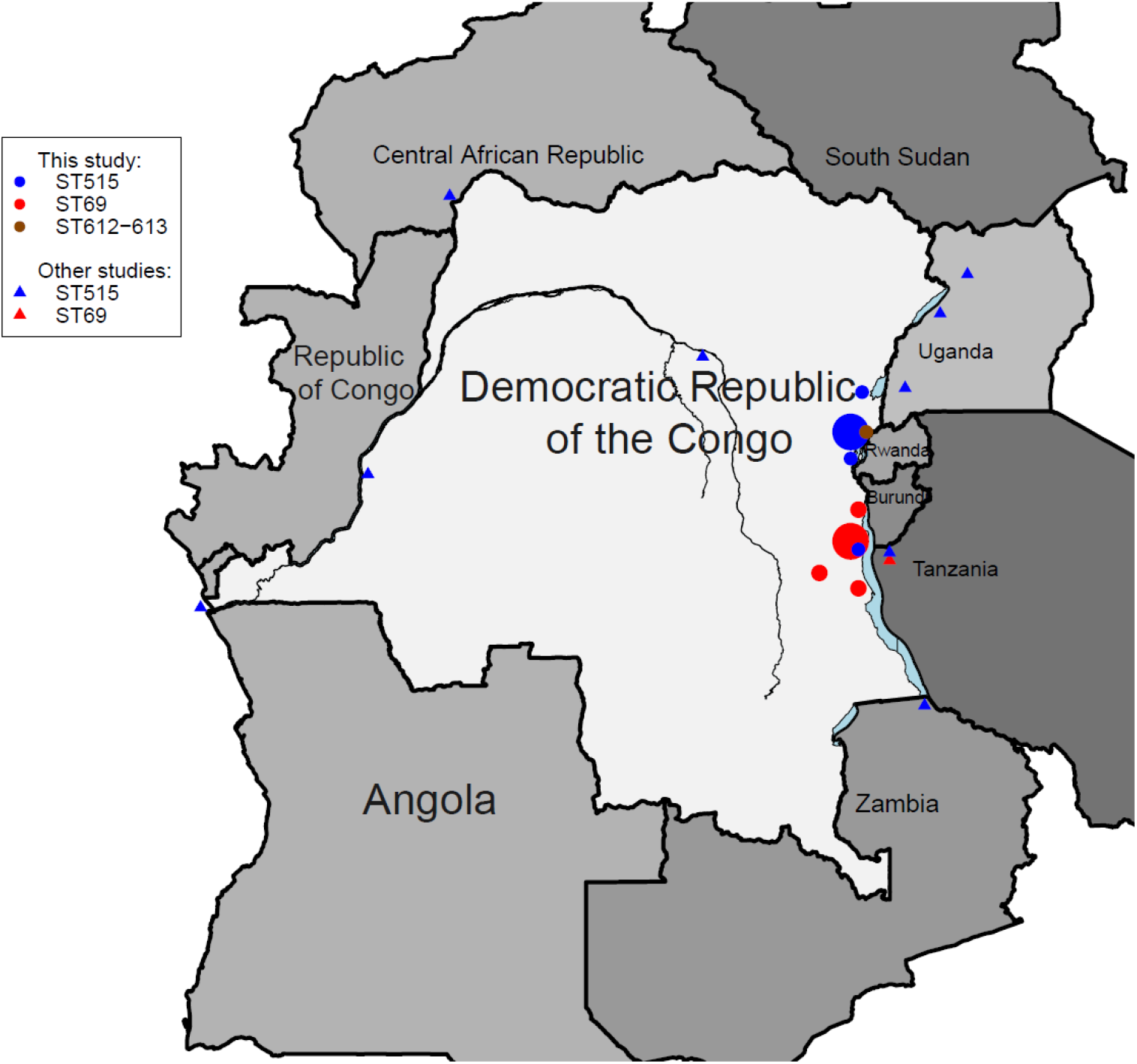
Geographical location of the sequenced *V. cholerae* isolates. This map was created using the Raster package [32], implemented in R statistical software version 3.6.1. The size of spots is somewhat correlated with the number of isolates from patients at the location.

Compared to closely-related DRC ST69 isolates ST515 isolates displayed a higher genetic diversity with core genomes separated by 0-25 and 0-77 SNPs (median: 6 and 14), respectively whereas the distance separating ST69 and ST515 core genomes from *V. cholerae* N16961 were 127-142 (median = 130) and 141-170 (median = 148) SNPs, respectively. There were major genetic differences between ST69 and ST515 sub-clades among which the ST515-specific 5-nucleotide (nt 24-28, TGTAC) frameshift deletion in the *webT* gene, not found in ST69 and creating a premature termination codon (Table 2).

**Table 2.**
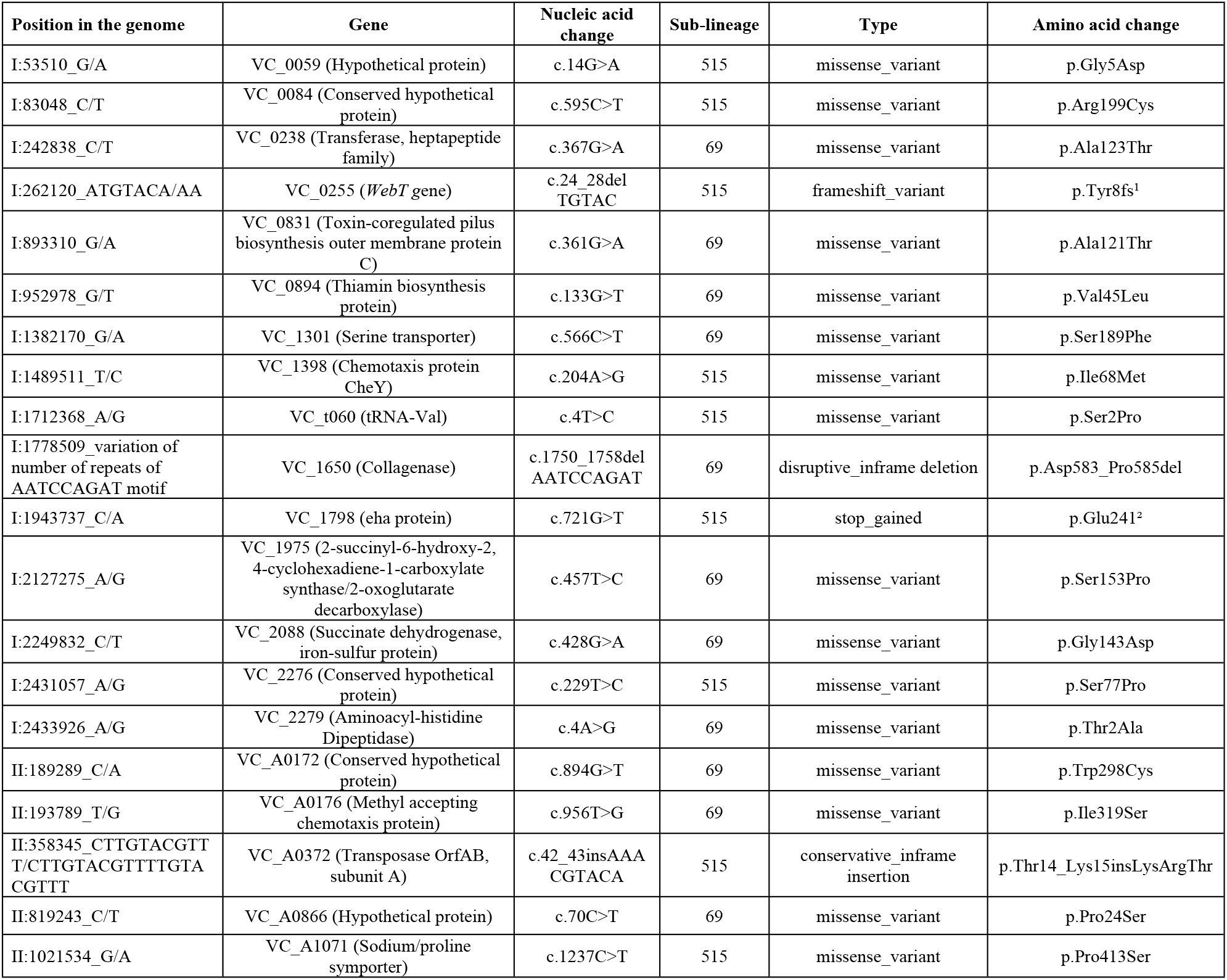
Major genetic differences between seven pandemic *V. cholerae* O1 sub-clades ST69 vs ST515 from eastern DRC. Only genetic changes impacting proteins are listed in the table

Legend: 1. Tyr8fs: frameshift after the 8th amino acid (tyrosine) in the webT protein; 2. Stop codon after the 241st amino acid (glutamate) in the VC_1798 (eha protein).

Other major discriminating genetic changes between the two sub-clades included (i) a variation of the 9-nt repeat AATCCAGAT corresponding to a DNP amino acid motif in the VC_1650 (chromosome I) of *V. cholerae* O1 isolates, with 6 versus 7 repeats for ST69 and ST515 sub-clades; respectively, (ii) the insertion of the AAACGTACA motif corresponding to KRT amino acids in the VC_A0372 (chromosome II), and (iii) the 721G→T transversion in the VC_1798 (chromosome I) leading to the apparition of a premature stop codon in the gene.

The 5 *V. cholerae* non-O1/non-O139 eastern DRC isolates did not cluster with representatives of *V. cholerae* O1 (Classical O1, Gulf Stream, pre-7^th^ and 7PET) (Fig. 3). These non-O1/non-O139 isolates, which are the first to be reported in DRC, did not carry any prophage associated with *V. cholerae*. They were assigned to two novel sequence types, i.e. ST 612 and ST 613, by the curator of the *V. cholerae* MLST database (https://pubmlst.org/vcholerae/). Whereas ST612 isolates (n = 4) were closely related between them and to some extent to isolates characterized in Mozambique [33] and Haiti [34], the ST613 DRC isolate could not be related to any characterized *V. cholerae* isolate.

**Fig. 3.**
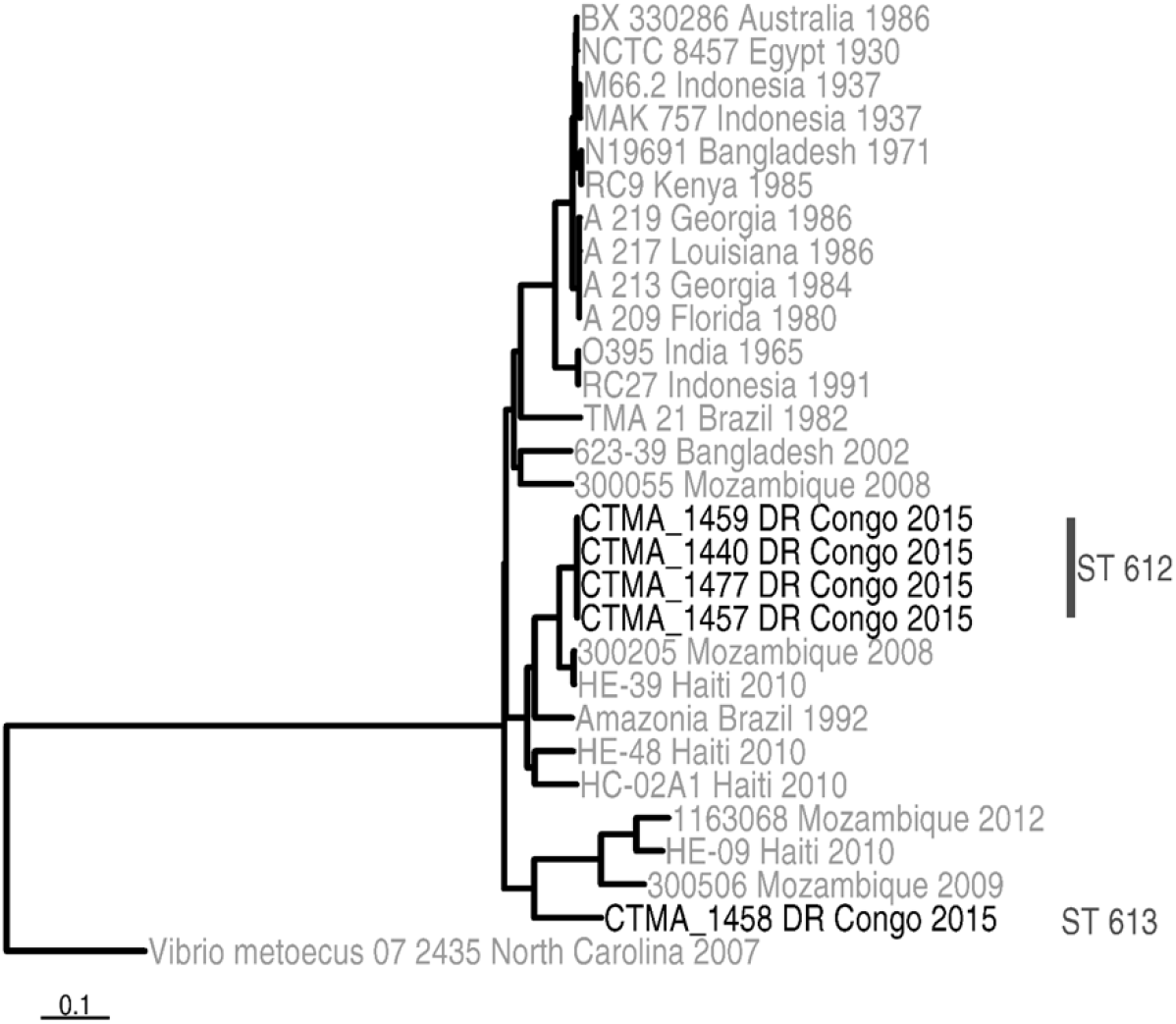
Phylogeny of the five *V. cholerae* non-O1/non-O139 from eastern DRC and their potential relationship with *V. cholerae* O1 and *V. cholerae* non-O1/non-O139 from other regions of the world.

With respect to the virulence genes, DRC O1 isolates harbored several virulence genes. Besides those associated with the CTX-3 prophage (*ctxA*, *ctxB*^Class^, *zot*, *ace* and *cep*), they all carried the following pathogenicity islands (PAIs): (i) the *Vibrio* pathogenicity island-1 (VPI-1), which harbors the genes encoding the toxin co-regulated pilus (TCP) and a cluster of four accessory colonization factor (*acfA, acfB, acfC* and *acfD)* genes [35], (ii) the *Vibrio* Seventh Pandemic Island-I (VSP-I), a 16-kb region which spans ORFs VC_0175 to VC_0185 and which is present only in O1 El Tor and O139 serogroup isolates [36], and (iii) the *Vibrio* Seventh Pandemic Island (VSP-II) with a large deletion spanning from ORF VC_0495 to VC_0512 deletion, as previously reported in several East African *V. cholerae* O1 isolates [22, 37]. In addition, DRC O1 isolates from this study harbored other virulence genes, among which the hemolysin A (*hly*A), the rtx (repeats in toxin) cluster (*rtx*A, *rtx*B, rtxC, rtxD genes), the virulence-associated (*vas*) operon, the *tox*R and *tox*T which activate numerous genes involved in virulence, as well as several genes of the type VI secretion system (T6SS), namely *hcp*, VCA0109, VCA0122, *vgrG.*2*, vgrG*.3*, vip*A and *vip*B genes [38]. However, these isolates lacked several virulence genes such as the WASA-1 [13, 39], *stn* and *NAG-S*. Whereas non-O1/non-O139 *V. cholerae* isolates from DRC lacked most of PAIs found in 7PET, they still harbored several virulence genes, namely members of the T6SS (VCA0109, VCA0122, *vgr*G.3), the *rtx* cluster (*rtx*A, *rtx*B, rtxC, *rtx*D genes), and the virulence-associated (*vas*) operon. The unique *V. cholerae* non-O1/non-O139 ST613 isolate lacked the *tox*R and *vgr*G.2 genes which were present in other four non-O1/non-O139 ST612 isolates.

Regarding the identification of antimicrobial resistance genes, all DRC O1 isolates (n=73) harbored the integrase gene of the SXT element (Int_SXT_) and the SXT/R391 integrative conjugative element (ICE) ICEVchBan5 which encodes resistance to several antibiotics, including sulfamethoxazole and trimethoprim [40, 41]. It is of note that ICEVchBan5 was lacking in non-O1/non-O139 isolates. In addition, all DRC O1 isolates harbored the APH3-DPRIME, APH6, drfA, dhfr, floR and SulI antimicrobial resistance genes. They also harbored the 248 G→A SNP in the quinolone-resistance determining region (QRDR) of the gyrA gene (VC_1258), resulting in the S83I substitution in the gyrA protein. All ST69 plus two DRC O1 ST515 (CTMA-1453 and CTMA-1454) isolates displayed the 254 G→A SNP in parC gene (VC_2430), resulting in the S85L substitution in that gene. No additional SNPs were found in the QRDRs of gyrA, gyrB, parC, and parE genes, nor were genetic determinants of beta-lactam resistance identified in these isolates. Among non-O1/non-O139 isolates, only ST612 harbored qnrVC and SulII genes. Conversely, ST613 was the only V. cholerae isolate to harbor the beta-lactamase carB gene.

## Discussion

In line with the phenotypic features [42], the WGS-based analysis of 78 *V. cholerae* isolates from eastern DR Congo (2014-2017) confirmed that all O1 (n=73) were 7PET variants (3^rd^ wave and T10 transmission event) genetically linked to the eastern African clade [5]. O1 isolates clustered closely in 2 distinct sub-clades consisting of ST69 and the newly assigned ST515. It is worth noting that the complete and/or draft genomes of ST515 were already available in public database but not assigned as ST515. The relatedness of O1 isolates within sub-lineage T10 was supported by SNP-based phylogeny and common genetic features among which the presence of CTX-3 prophage, VSP-II and several AMR genes, and a lack of WASA-1 in line with previous characterization of *V. cholerae* isolates from DRC collected during the period 2006-2014 [5, 43]. In all O1 eastern DRC isolates, a consistent low susceptibility to nalidixic acid without resistance to ciprofloxacin was correlated with the presence of the S83I substitution in *gyr*A. Moreover, a S85L substitution in *parC* was found in all ST69 isolates and two ST515. Interestingly, a recent study on *V. cholerae* isolates associated with cholera outbreaks in Yemen linked the presence of *gyr*A (S83I) and *par*C (S85L) substitutions with a decreased susceptibility to ciprofloxacin [44]. However, current *V. cholerae* isolates from eastern DRC differed from those from Yemen as only 6 out of 39 ST69 eastern DRC isolates carrying both *gyrA* (S83I) and *ParC* (S85L) substitutions actually showed a reduced susceptibility to ciprofloxacin, and this observation was in agreement with previous data [45].

Unlike other African countries where further introduction events (i.e. T11, T12 and T13) within the 3^rd^ wave of the 7PET have been reported [15], it is noteworthy that isolates from eastern DRC all belonged only to the T10 introduction event. These results suggest that these T10 isolates have firmly established themselves in the Congolese Albertine rift, becoming an autonomous source of endemic, sporadic and epidemic cholera in the eastern DRC sub-region.

Several genetic features differentiated *V. cholerae* O1 ST69 and ST515 sub-clades from eastern DRC (Table 2), highlighting the continuous local evolution and adaptation of O1 isolates and supposedly determining their particular geographical distribution pattern. This adaptive potential might indeed be triggered by changing environmental conditions, e.g. altitude, temperature, humidity and anthropogenic impacts, which, in turn, could potentially affect the interaction between the bacterium and its host. For instance, a variation in the number of ATAATCCAG motif repeat can affect *V. cholerae* growth depending on the range of incubation temperature [46]. Likewise, the serotype switch from Ogawa to Inaba in all ST515 isolates probably results from the *webT* gene inactivation consecutive to the 5-nt frameshift deletion as suggested earlier [47, 48]. This serotype switch could affect patient’s immune response to cholera in regions where O1 serotype Ogawa was predominant [47].

The *V. cholerae* O1 global phylogeny including data from Uganda [15], Tanzania [49], DRC, Central African Republic, and Zambia [5] confirms that the ST515 sub-clade has now spread to several regions of Central and Eastern Africa, including western provinces of DRC up to the Atlantic coast. Whereas the reason why only the ST515 expands so widely, and not the ST69 sub-clade, remains unknown, the hypothesis is that the higher genetic diversity among ST515 isolates results from a high mutation frequency, which favors their adaptation to changing environmental conditions.

As also reported in other countries [33,50,51], two *V. cholerae* non-O1/non-O139 lineages were also identified and characterized from cholera-like diarrhea cases in eastern DRC, and were assigned to two novel sequence types, ST612 and ST613. Recent cholera outbreaks affecting the Kasai provinces highlight the urgent need to better understand the factors favoring the endemicity and epidemicity of cholera among the exposed populations. As illustrated with ST69 and ST515 in this study, phylogenetic changes may be associated with local adaptation to eastern DR Congo, clonal expansion of *V. cholerae* sub-lineages and consecutive spread in neighboring DRC provinces and bordering countries.

## Acknowledgements

We acknowledge the assistance of Drs Chirimwami Marie-Paul and Kakule Michel (Minova healthcare zone, South Kivu province, DR Congo) for supporting local sample collection. We also thank Michèle Bouyer (Defense Laboratories Department) for providing assistance in culturing isolates, and the technical staff of AMI-LABO and of Centre de Diagnostic et de Recherche en Maladies for handling isolates. Finally, the authors wish to express their gratitude to the EMBL-EBI team (Genome Campus, Hinxton, Cambridgeshire, CB10 1SD, UK) for providing ENA accession numbers, and to Dr Sophie Octavia, the *Vibrio cholerae* MLST database curator, for the assignment of the MLST ST.

## S1 appendix

Quality metrics of genome assemblies

